# Soil amendments and suppression of *Phytophthora* root rot in avocado (*Persea indica*)

**DOI:** 10.1101/2022.01.31.478582

**Authors:** Qurrat Ul Ain Farooq, Jen McComb, Giles StJ. Hardy, Treena Burgess

## Abstract

The ability of microbial or mineral-based soil additives to suppress root rot caused by *Phytophthora cinnamomi* was assessed. Phosphite and metalaxyl treatments for the control of disease, and glyphosate for weed control were also assessed. A treatment simulating avocado orchard conditions had chicken manure, wood mulch, and mulch from beneath trees in an avocado orchard added to the pots. Soil treatments (three probiotic and two mineral-based) were applied to 9-month-old saplings growing in containers in a glasshouse. After one-month, half of the plants of each treatment were inoculated with the pathogen. Three months after inoculation, plants were harvested and plant growth and root damage were measured. In the first experiment infestation with *P. cinnamomi* significantly reduced fine root dry weight in all plants except those in soil treated with one silicon-based mineral mulch. Visible root damage was higher in plants treated with probiotics. In this experiment, and in a repeat experiment the reduction of fine root damage achieved by spraying plants with phosphite or addition of a silica based mineral mulch was similar. Phosphite was preferable to metalaxyl as a chemical treatment, as the latter reduced shoot and root growth of non-infected plants. Glyphosate treatment of wheat seedlings growing in the pots with the avocados reduced shoot and fine root growth of both non-infected and infected plants. These observations need to be confirmed under field conditions.

## Introduction

Phytophthora root rot caused by the oomycete *Phytophthora cinnamomi* is one of the worst avocado diseases worldwide. This pathogen attacks the feeder roots of avocado trees, and if left untreated can result in the death of trees and high economic losses (Reeksting et al. 2016; Ramirez□Gil et al. 2017). Management strategies include controlling movement of people, livestock and vehicles, the use of chemicals, soil additives and development of *P. cinnamomi*-tolerant cultivars or rootstocks (D’Souza et al. 2005). The most robust and environmentally benign methods are integrated management approaches (Pegg and Whiley 1987; Wolstenholme and Sheard 2010) including selection of tolerant varieties, application of organic fertilizers and mulches, addition of cultured microbial antagonistic agents, inorganic nutrition and liming, irrigation management and fungicide application (Wolstenholme and Sheard 2010; Pegg 2010). Phosphite is the most commonly used chemical for control of *Phytophthora* root rot caused by *P. cinnamomi* in agricultural, forestry, horticultural and natural ecosystems worldwide (Pegg et al. 1987; Hardy et al. 2001; Shearer and Fairman 2007; King et al. 2010; Scott et al. 2015; Ramírez-Gil et al. 2017; Masikane et al. 2020). It is the only chemical effective against this disease as it acts directly on the pathogen as well as increasing the plants’ resistance to the pathogen (King et al. 2010). However, prolonged use of phosphite increases the potential for development of phosphite resistance in the pathogen (Dobrowolski et al. 2008). Moreover, use of agrochemicals may have a negative effect on environment through detrimental effects on beneficial soil microbes, and disruption of soil ecology (Hardy et al. 2001; Gill and Garg 2014; Xi et al. 2020).

Commercial microbial soil probiotic additives/conditioners comprising proprietary mixtures of microbial species are receiving increasing attention with the global demand for these products growing at 10% each year (Berg 2009; Song et al. 2012). The most promising biocontrol beneficial microbial organisms, effective against several foliar and soil-borne diseases are the Proteobacteria (*Bacillus* spp.), actinobacteria (*Streptomyces* spp.), the fluorescent pseudomonads (e.g. *Firmicutes*), and fungi (e.g. non-pathogenic *Fusarium* spp. and *Trichoderma* spp.) (Raaijmakers et al. 2009; Bhattacharjee and Dey 2014).

Mineral soil conditioners, particularly silicon-based mulches, have also shown encouraging outcomes for plant disease control and growth in agricultural crops (Pozza et al. 2015; Tubana et al. 2016). They have been effective for the control of anthracnose (*Colletotrichum lindemuthianum*) (Moraes et al. 2009) and powdery mildew (*Sphaerotheca fuliginea*) (Samuels et al. 1991; Menzies et al. 1992; Belanger et al. 2003). Silica is known to impact on a wide range of plant metabolic processes, and also to affect the soil microbiota, though less is known about the latter (Rajput et al. 2021).

No information is currently available on the comparative effectiveness of soil probiotics developed commercially for management of *Phytophthora* in avocado. This study presents information on the effectiveness spraying with phosphite, compared with use of organic mulches, commercial soil probiotics and silicon-based mineral mulches on root damage in avocado caused by *P. cinnamomi*. It also investigates whether a commonly used herbicide, glyphosate may exacerbate Phytophthora root damage.

## Methods

### Plant Material

Avocados (cv. Reed) were grown from seed, initially in potting mix (Richgro Garden Products, Australia). Nine-month-old plants were transplanted into 220 x 330 mm (15 L) free-draining polybags (Garden City Plastics, Forrestfield, Western Australia) for the first experiment, and for second experiment 150 x 380 mm (7 L) polybags were used.

### Soil and growing conditions

In experiment 1 Channybearup yellow brown clay loam soil was sourced from Manjimup Western Australia (WA). It is moderately well draining and has a high-water holding capacity. It was used in a 1:4 ratio of soil: coarse perlite (Perlite and Vermiculite Company, Myaree, WA). For the repeat experiment, a clay loamy soil was collected from Carabooda, Western Australia (WA). It was moderately well drained with good porosity which was mixed with river sand in ratio of 1:1. Both soils are known to be conducive to *Phytophthora* as native vegetation or avocado orchards on these soils suffer from phytophthora dieback but both soils were determined to be free of *P. cinnamomi* through standard baiting of the soil (Aghighi et al. 2016; Simamora et al. 2018). Two inoculation tubes (25 mm dia x 20 cm long) were placed in each bag when plants were potted to minimise the chance of damaging roots during the insertion of the inoculation plugs. Plants were watered daily to container capacity. Plants were placed on benches in a completely randomized design in evaporative cooled glasshouse maintained at 25-27 °C. All had weekly applications of liquid Thrive for fruits (Yates, Australia), (4gm/L, 300-400ml/pot) and 5ml/L Eco oil (Organic crop protectant Pty. Ltd., NSW, Australia) was sprayed on foliage and the soil surface when required for insect control.

### Treatments

There were eleven treatments in Experiment 1, and treatments 1, 3, 4, 7, 8,10 were repeated in experiment 2.

Treatment 1 had no additives.

Treatments 2-11 had 50 g of well decomposed chicken manure applied monthly and 200 g/pot jarrah (*Eucalyptus marginata*) wood mulch placed on the surface at the time of transplanting. In experiment 2, in the smaller pots, half these amounts were applied.

Treatments 3-11 had 150 g of avocado mulch added at the time of transplanting for experiment 1, (75g was applied in experiment 2). Avocado orchard mulch was collected from a 20-year-old orchard in which *P. cinnamomi* is controlled by phosphite spray (Delroy Orchards, Pemberton, WA), application of decomposed wood mulch, chicken manure, fertigation with standard fertilisers and fish emulsion. The jarrah wood mulch and the avocado orchard mulch were determined to be free of *P. cinnamomi* through baiting.

Treatment 4 had weekly application of a 25% concentration of a microbial soil conditioner known to include *Lactobacillis, Bacillus, Saccharomyces, Acetobacter* and *Azotobacter*.

Treatment 5 had an additive (conc. 1:500) sprayed to foliage to run-off and applied to soil to field capacity (immediately after potting, and after inoculation). This additive contained humates and an organic chemical.

Treatment 6 had applications of two additives that contained a mixture of *Bacillus* species. One was applied at 50 g/pot at the time of transplanting, and the second at 2.5 ml/pot every 6 weeks.

Treatments 7 had 200 g/pot for experiment 1 and 100 g/pot for experiment 2 of ‘Mineral Mulch’, a mulch containing calcium, silicon (www.mineralmulch.com)

Treatment 8 had applications of mineral based additives containing silica and sulphur. At the time of transplanting 2 g/pot of Dolomite (calcium magnesium carbonate) and 1.5 g/pot of Humus 400’ (derived from. lignite; for inorganic analysis available: ecogrowth.com.au/products/humus400) was added. Then every fortnight 5 g/pot of ‘Eco-prime’ (chemical analysis available: ecogrowth.com.au/products/eco-prime-avocado) and 0.5 ml of ‘Eco X’ (containing silica, calcium and sulphur) was added for both experiments.

Treatment 9 had wheat seeds planted at the time of transplanting. When four weeks old the wheat seedlings were sprayed with ‘Roundup’ (active ingredients: 7.2g/L glyphosphate and 21g/L Nonanoic acid manufactured by Evergreen Garden Care Australia Pty. Ltd., NSW, Australia) to run-off) (the stems of the avocados were protected with aluminium foil). In avocado orchards glyphosate is routinely used to control weeds (Nartvaranant et al. 2004).

Treatments 10 had plants sprayed with fungicide for *P. cinnamomi* control:Agri-Fos 600 (active ingredient: 600g/L phosphorus acid manufactured by Nufarm Australia Ltd, Western Australia) sprayed according to manufacturer manual on the avocado plants to runoff. Phosphite was applied after transplanting and repeated after a further 10 days.

Treatment 11 was also a fungicide treatment: Ridomil Gold (active ingredient: 25g/kg Metalaxyl-M; Syngenta Australia Pty Ltd, NSW, Australia) was applied as a soil drench at the recommended rate. Metalaxyl was applied 2 weeks before plants were inoculated with *Phytophthora*, then 6 weeks after inoculation.

### Inoculum production and inoculation

Branches of live tagasaste (*Chamaecytisus palmensis*) (1-2 cm dia.) were cut into approximately 2 cm pieces. Two hundred plugs were placed in each of ten 2 L conical flasks. The plugs were soaked in distilled water overnight, then rinsed. Distilled water (approx. 50-70 mL, sufficient to cover the bottom of the flask) was added, the flask was plugged with a non-absorbent cotton plug and autoclaved at 121 °C for 30 min, then allowed to cool to room temperature and the process was repeated after 24 h. Two Petri plates of *P. cinnamomi* (isolate MP 94-48 from the Centre for Phytophthora Science and Management culture collection, Genbank Accession number for ITS gene region is JX113294), cultured on a V8 (vegetable juice) agar medium for 7 days at 25 °C in the dark, were cut into 1 cm squares and aseptically added to the conical flasks. The flasks were shaken to evenly distribute the agar plugs and incubated at 25 °C in the dark. Flasks were incubated for 4 weeks and shaken weekly to obtain uniform colonisation of the inoculum plugs.

All treatments consisted of 10 non-inoculated controls and 10 infested plants inoculated with *P. cinnamomi*. Plants were inoculated by insertion of two *P. cinnamomi* colonised plugs to a depth of 10 cm into each of the two holes created after removal of the inoculation tubes. Similarly, for the control plants, non-colonised plugs were inserted to the same depth in the holes after removal of tubes in non-inoculated pots. Three weeks after inoculation, all the plant containers were flooded for 24 h to stimulate the release of zoospores.

### Harvest

Twelve weeks after inoculation with *P. cinnamomi* the plants were harvested. In both experiments, shoots were excised then dried in an oven at 60 °C to constant dry weight. Roots were washed free of soil and the visual damage caused by the pathogen scored using the rankings 1= healthy roots, 2= 1-25% damage, 3= 26-50% damage, 4= 51-75% damage and 5= 76-100% damage. Fine roots were separated, and the dry weight of fine and coarse roots recorded. Total root dry weights were calculated by adding fine and coarse root dry weights. Re-isolation of *P. cinnamomi* was undertaken using a randomly chosen plant in each treatment to confirm that the inoculation caused disease.

### Data analysis

To check the effect of different treatments on the control of damage caused by *P. cinnamomi*, data for shoot dry weight, total root dry weight and fine root dry weight, these data were analysed using a linear model with terms for treatment, + and - *P. cinnamomi*, and their interactions. Homoscedasticity was assessed by plotting residuals vs fitted values and normality of residuals were assessed using q-q plots. Duncan multiples range test was used to assess the significance between all pairs of treatments /+,- *P. cinnamomi* combinations. The qualitative data on root damage for each treatment was assessed separately using Chi square tests. All the data analyses were conducted and bar graphs were generated in R software.

## Results

### Shoot dry weight

In both experiments all seedlings were alive at the time of harvest and there was no evidence of wilt associated with the presence of *Phytophthora*. The first experiment for non-infested plants highest total shoot dry weight was recorded for plants receiving phosphite (T10) and this was statistically significantly higher than values for treatments with metalaxyl (T11) or glyphosate (T9) (Fig 2a). Following infestation with *Phytophthora* only plants treated with phosphite showed a significant fall in shoot dry weight (Fig 2a).

**Fig1.**
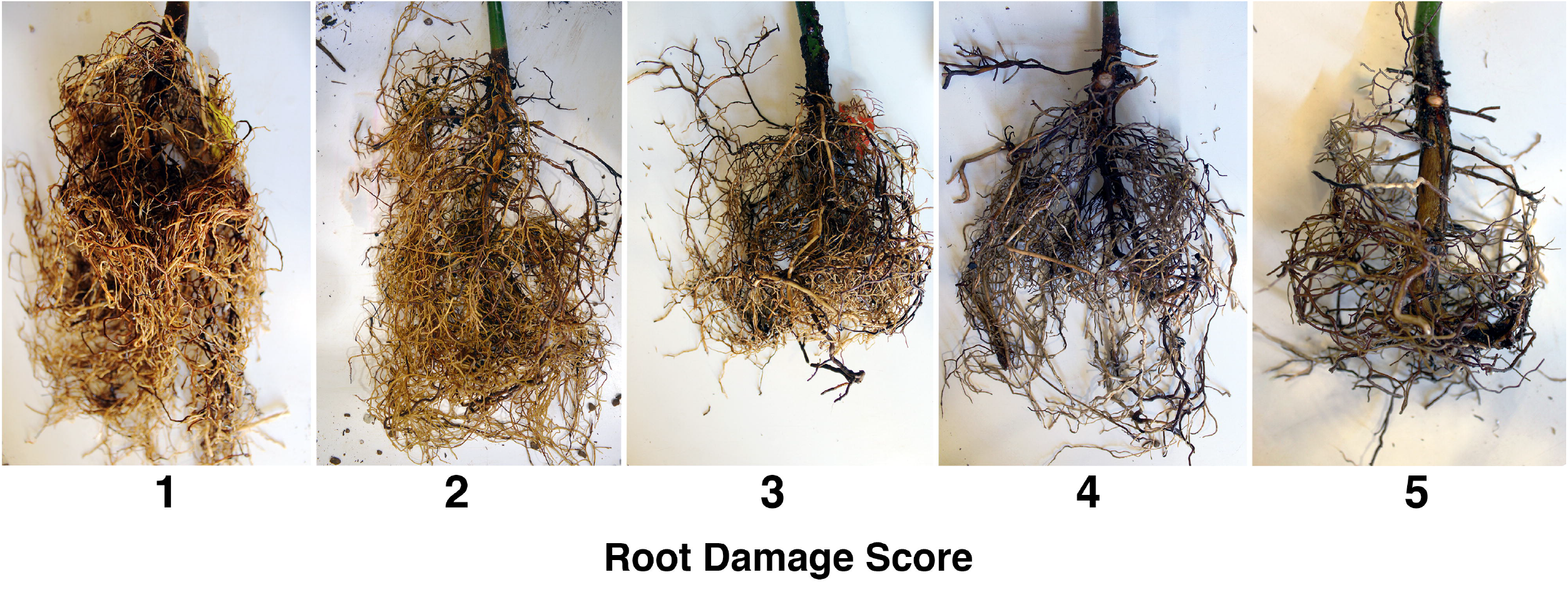
Examples of root damage of avocado roots after inoculation with Phytophthora cinnamomi. Whole root systems were rated visually for Phytophthora root rot on the bases of scale 1-5 (left to right) 1 = healthy roots, 2 = 1-25% damage, 3 = 26-50% damage, 4 = 51-75% damage 5 = 76-100% damage.

**Fig. 2.**
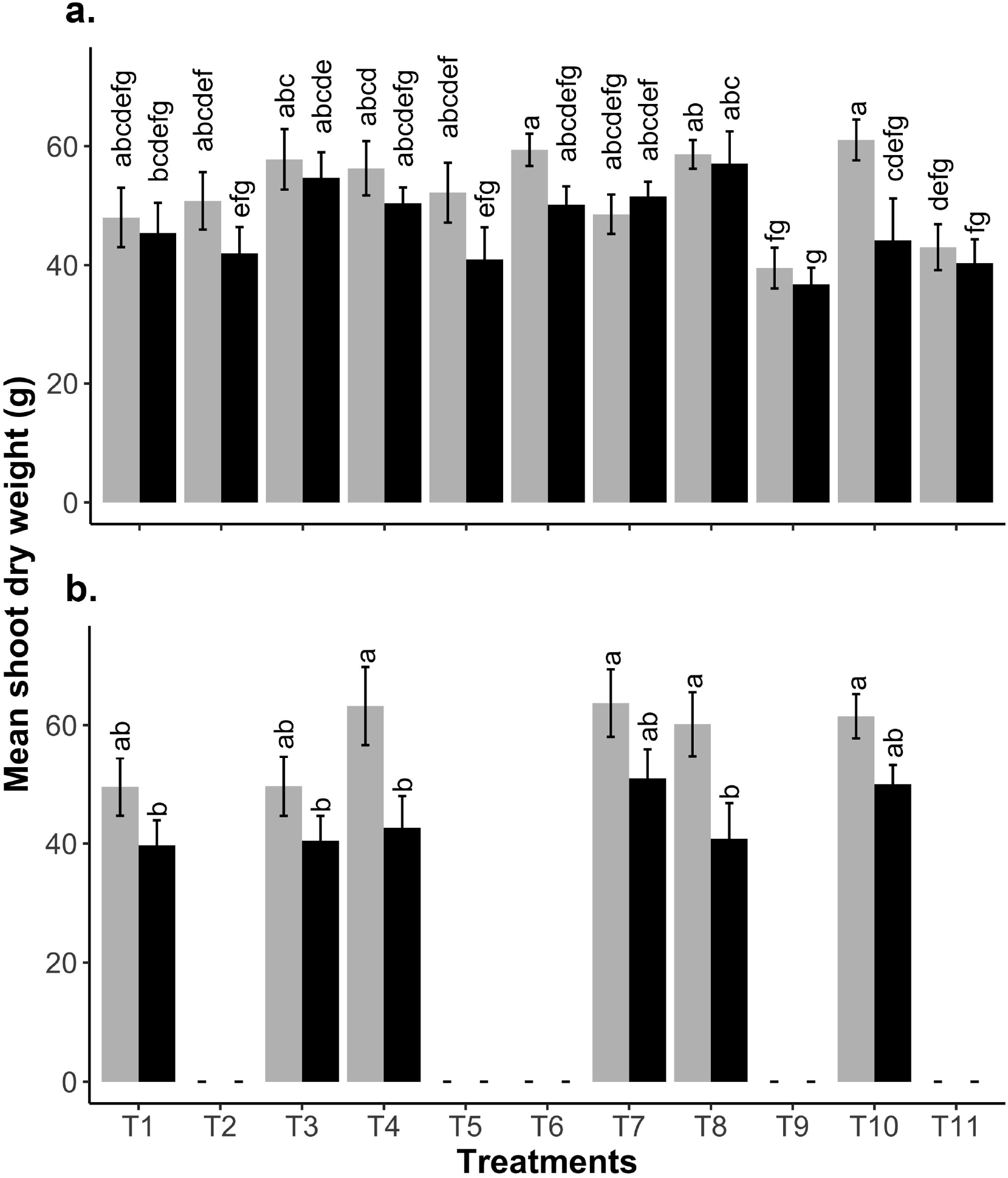
a) The effect of eleven soil treatments (T1-T11, see Table 1) on shoot dry weight of non-infested avocado plants (grey bars) and those infested with *Phytophthora cinnamomi* (black bars). b) The effect of soil treatments (T1, T3, T4, T7, T8, T10, see Table 1) on shoot dry weight of non-infested avocado plants (grey bars) and those infested with *Phytophthora cinnamomi* (black bars). Vertical lines indicate SE and letters indicate significant difference (p< 0.05).

When selected treatments were repeated, there was no significant difference in shoot dry weight of treatments not infested with *Phytophthora*, while in infested plants shoot dry weight was highest for plants treated with mineral mulch 1 (T7) or phosphite (T10), but there was no statistical difference between these weights and the other treatments (Fig 2b).

There was a significant reduction of shoot dry weight following infection in the treatments with probiotic 1 (T4) or mineral mulch 2 (T8) (Fig 2a).

### Total root dry weight

In the first experiment for non-infested plants highest total root weight was recorded for plants receiving mineral mulch 2 (T8) but this was only statistically significantly higher than values for treatments with mineral mulch 1 (T7) metalaxyl (T11) or glyphosate (T9) (Fig 3a). In the repeat experiment total root dry weight in non-infested plants was significantly higher in treatments with probiotic 1 (T4), mineral mulch 1 (T7) and phosphite (T10) than in the treatment with avocado mulch or with no additives (Fig 3b). When soil was infested with *Phytophthora* in the first experiment there was a significant drop in total root weight for the treatment with no additives or with only chicken manure, but no significant reduction in the other treatments (Fig 3a).

**Fig. 3.**
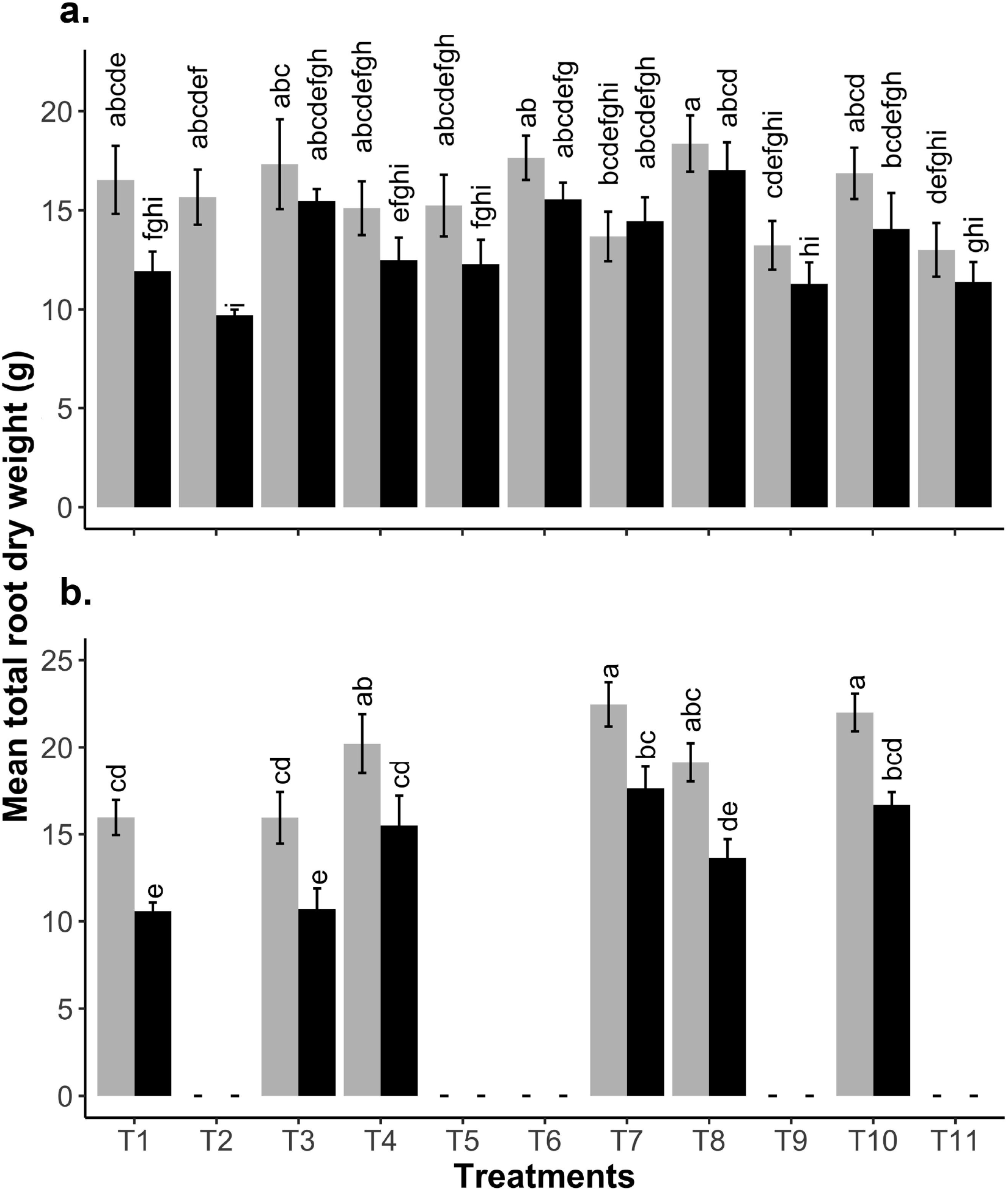
a) The effect of eleven soil treatments (T1-T11, see Table 1) on total root dry weight of non-infested avocado plants (grey bars) and those infested with *Phytophthora cinnamomi* (black bars). b) The effect of soil treatments (T1, T3, T4, T7, T8, T10, see Table 1) on total root dry weight of non-infested avocado plants (grey bars) and those infested with *Phytophthora cinnamomi* (black bars). Vertical lines indicate SE and letters indicate significant difference (p< 0.05).

In repeat experiment, in both non-infested, and infested soil, probiotic 1, mineral mulch and phosphite showed more root growth with greatest growth being observed in plants from the mineral mulch 1 and phosphite treatments. However, the reduction in root dry weight after infection was significant in all treatments (Fig 3b).

### Fine root dry weight

For the non-infested treatments, fine root dry weight was not significantly increased by the additional of any soil additives (T4-T8) above that of adding avocado orchard mulch alone (T3) (Fig. 4a). For non-infested plants, phosphite (T10) also had no impact on the fine root dry weight, however, metalaxyl (T11) and glyphosate (T9) both resulted in a significant reduction in the fine root dry weight (Fig 4a). The fine root dry weight was reduced by infection with *P. cinnamomi* in all treatments except T7 (mineral mulch1) (Fig. 4a). This reduction was significant in the treatments with no mulch, chicken manure (T2), organic mulch (T3), T4 (probiotic conditioner 1) and T5 (probiotic conditioner 2) (Fig 4a). The result was not significant for any of the other treatment pairs suggesting that the other additives (T6, T7 and T8, T9, T10, T11) reduced the impact of *Phytophthora* on the loss of fine roots. This was most striking for mineral mulch 1 (T7), where the fine root weight of infested and non-infested seedlings was the same (Fig. 4a).

**Fig. 4.**
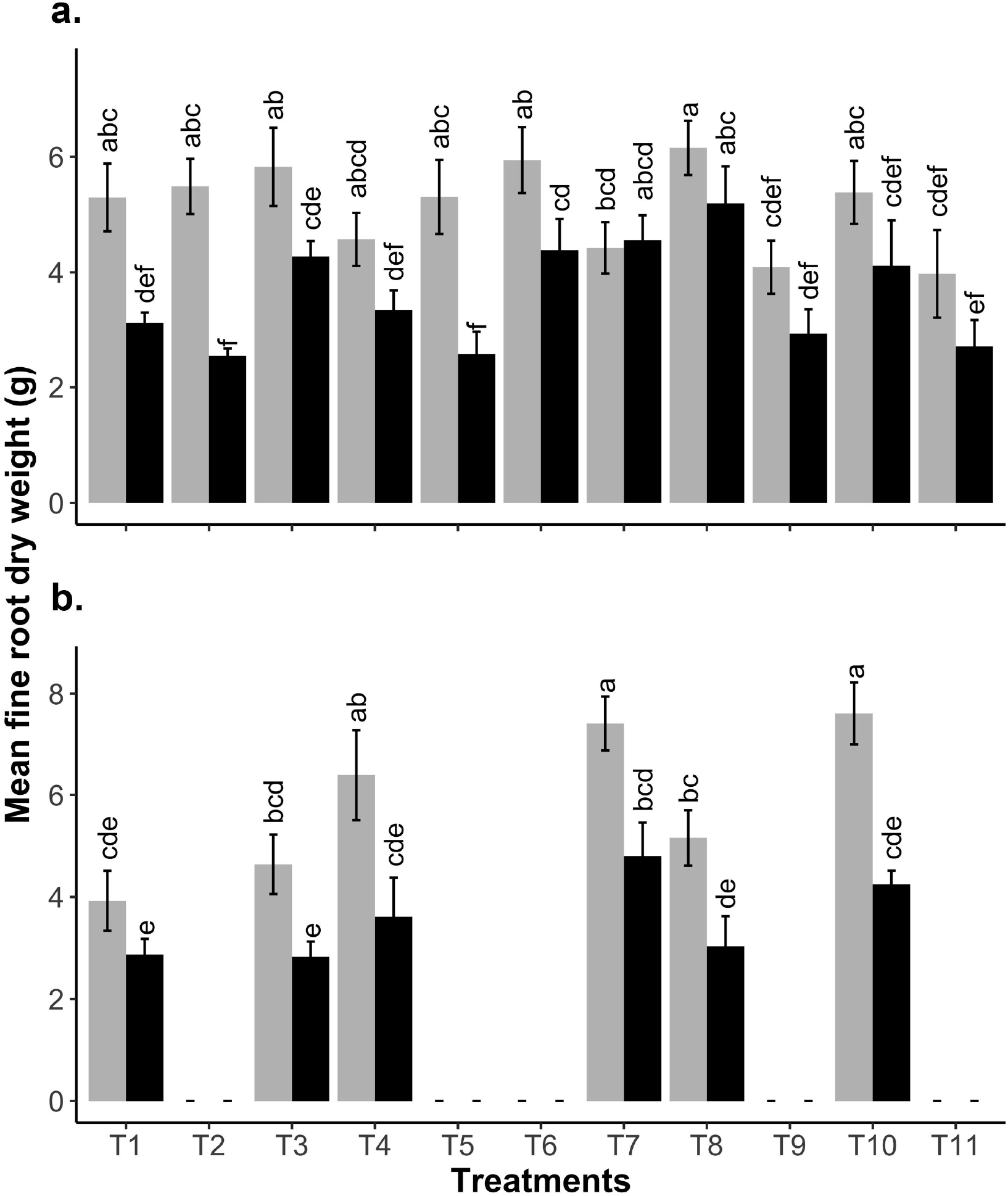
a) The effect of eleven soil treatments (T1-T11, see Table 1) on fine root dry weight of non-infested avocado plants (grey bars) and those infested with *Phytophthora cinnamomi* (black bars). b) The effect of soil treatments (T1, T3, T4, T7, T8, T10, see Table 1) on fine root dry weight of non-infested avocado plants (grey bars) and those infested with *Phytophthora cinnamomi* (black bars). Vertical lines indicate SE and letters indicate significant difference (p< 0.05).

In the repeat experiment in non-infested soil, application of mineral mulch 1 (T7) or phosphite (T10) gave the highest fine root dry weight. For plants grown in infested soil, treatment with mineral mulch 1(T7) or probiotic conditioner 1(T4) gave results equivalent to treatment with phosphite with the highest fine root weight being recorded for plants treatment with mineral mulch 1 (T7) (Fig 4b). However, after infection significant reduction in fine root dry weight was observed in all other treatments except no mulch (T1) (Fig 4b).

### Root damage

There was no significant difference in the root damage rating between treatments for non-infested plants but significantly (P<0.05) more root damage in all treatments infested with *P. cinnamomi* compared to their respective non-infested controls (Chi square test) in both experiments (Figs. 5a, b). In experiment 1 for infested plants there was greater root damage in treatments with probiotic conditioners (T4, 5, 6) and the glyphosate treatment. Infested plants given T8 (Ecoprime, a silicon-based mineral conditioner) had less damage than those in T3 (avocado orchard mulch) while in mineral mulch (T7) damage of infested plants was similar to that seen in avocado orchard mulch (T3). Amongst the pesticide treatments, glyphosate application (T9) resulted in more root damage of infested plants than infested plants treated with phosphite (T10) and metalaxyl (T11). Application of avocado mulch (T3), or mineral mulch (T7, T8) resulted in resulted in similar or less damage than that seen on plants sprayed with phosphite (T10) (Fig 5a).

**Fig. 5.**
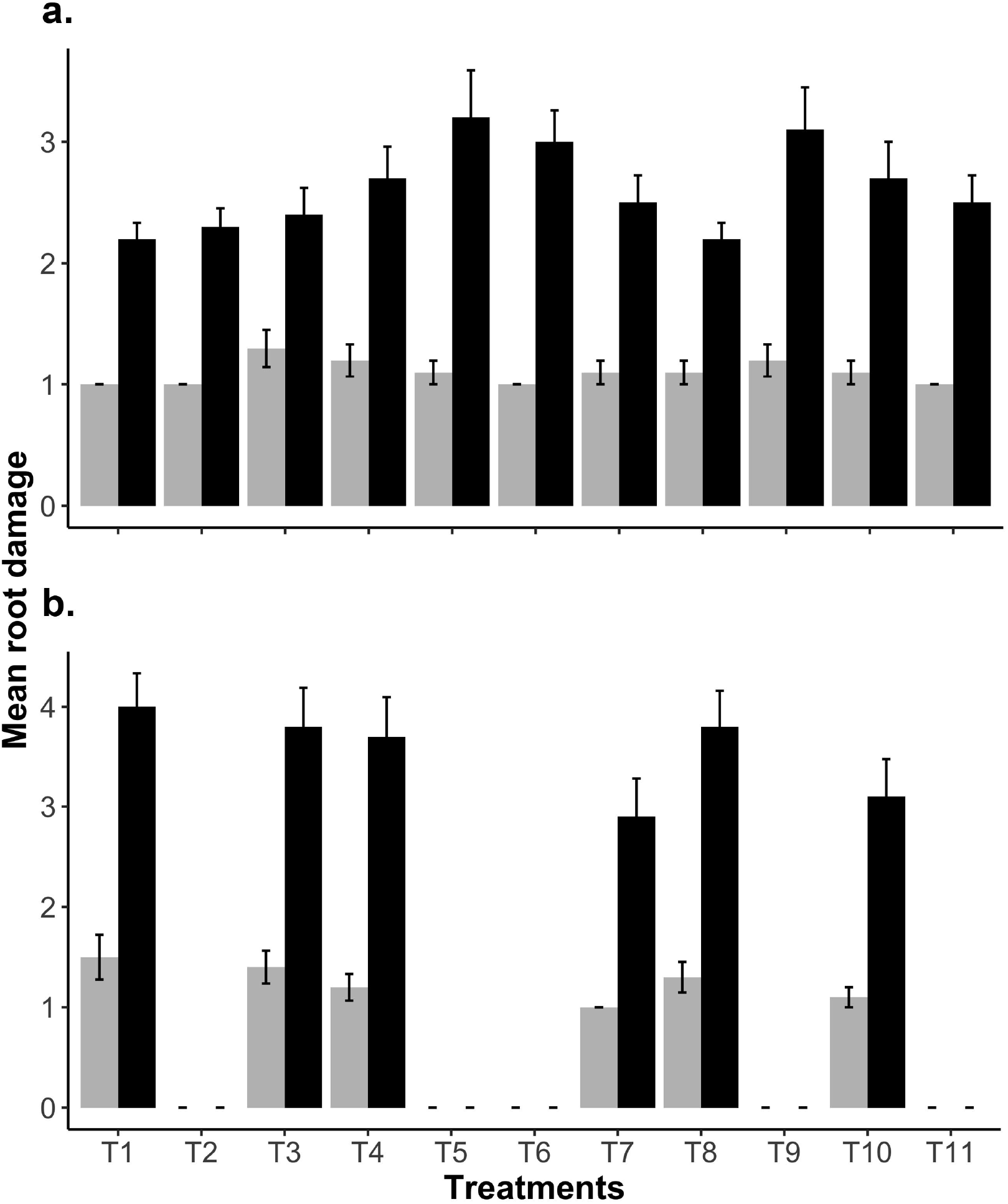
a) The effect of eleven soil treatments (T1-T11, see Table 1) on root damage of non-infested avocado plants (grey bars) and those infested with *Phytophthora cinnamomi* (black bars). b) The effect of soil treatments (T1, T3, T4, T7, T8, T10, see Table 1) on root damage of non-infested avocado plants (grey bars) and those infested with *Phytophthora cinnamomi* (black bars). Vertical lines indicate SE and letters indicate significant difference (α< 0.05).

In the second experiment overall root damage was higher in both non infested and infested treatments than in experiment 1. In infested soil the least root damage was in plants with mineral mulch 1 or phosphite. This reduction was significantly less compared to plants with no mulch (T1), avocado orchard much (T3) or mineral mulch 2 (T8) (Fig 5 b).

## Discussion

In these experiments although fine root weight was reduced by up to 50% in infested plants in some treatments, and 51-75% of the roots showed visible damage, shoots were healthy and no wilting was observed. This is probably because plants were grown under glasshouse conditions with adequate daily watering. Under field conditions, similar root damage could be expected to have much greater impact on total dry weight. In addition, if the glasshouse experiment had utilized younger plants with less woody roots (Rodriguez□Molina et al. 2002), or if the plants had been allowed to grow for longer after inoculation (Faber et al. 2000), there may have been a greater impact of *P. cinnamomi* on total dry weights and more divergence in effectiveness of the different treatments as assessed by total dry weight. As *P. cinnamomi* has its initial impact on fine roots, it is considered that the most accurate assessment of the effectiveness of the treatments applied here is seen through the data on total root dry weight, fine root dry weight and root damage.

### Effect of treatments on plants in soil infested with *P. cinnamomi*

The addition of avocado orchard mulch (T3) resulted in a small improvement in root dry weight compared to those without (T1 and T2) in experiment 1 but not in the repeat experiment. Treatment 3 is the closest equivalent to conditions in an avocado orchard in these pot experiments. No consistent beneficial effects of other soil additives on total root dry weight, fine root dry weight and root damage were seen. A possible exception is mineral mulch (T7), for which total root dry weight and fine root dry weight was not reduced by *P. cinnamomi* and there was less visible root damage than in other treatments. In the second experiment avocado mulch gave no improvement, but addition of probiotic 1 (T4), mineral mulch (T7) or phosphite (T10) increased both root total weight and fine root weight. Visible root damage was increased by addition of treatments to the avocado mulch (T3) treatment except for mineral mulch (T7). In experiment 2 mineral mulch (T7) showed the lowest damage. Mineral mulch 1 (T7) contains silicon, which is known to improve root growth and increase defence mechanisms of roots when infected with *Phytophthora* (Bekker et al. 2006; Bekker et al. 2007; Bekker 2011; Dann and Le 2017).

### Effect of the treatments on roots of non-infected plants

In experiment 1 no treatment significantly affected total root dry weight, fine root dry weight, or visible rot damage in any non-infested treatment. In the second experiment total root dry weight and fine root dry weight was increased by application of probiotic (T4), mineral mulch (T7 & T8), or spraying plants with phosphite (T10). Visible root damage was similar in all treatments. The soils were very similar in the two experiments but use of smaller pots, and sand rather than perlite in the mix for experiment 2, resulted in a greater root mass in the non-infested control treatment with no additives (T1) than in experiment 1, and conditions more conducive for *Phytophthora* as seen by the higher level of fine root damage. In experiment 2 the conditions were more suitable for detection of the effects of additional soil additives on root growth and reduction of Phytophthora root damage.

The addition of mineral mulch and phosphite had positive effect on the growth of non-infected plants and improved the root biomass. The silicon-based mineral mulch may act as fertilizer and improve the availability of micronutrients and cation exchange activity, and reduce the excess uptake of Fe, Mn and Al (Etesami and Jeong 2018; Etesami 2018) and thus improve growth of shoots and roots (Al-Garni et al. 2019). Phosphite may also act as fertilizer and stimulate the root growth (Ramirez□Gil et al. 2017).

### The effect of fungicides and an herbicide

In the current experiment it appeared that glyphosate use may worsen disease symptoms. (T11). Although no glyphosate was sprayed directly on the avocado plants, (it was used to kill a lawn of wheat growing in the pots) it reduced both total and fine root dry weight and there was a high level of root damage when *P. cinnamomi* was present (T11). The use of glyphosate in orchards may be damaging to avocado trees, possibly as it may result in mineral deficiencies which in turn increase disease susceptibility (Kremer et al. 2009; Zobiole et al. 2010; Zobiole et al. 2011). It may also be detrimental to beneficial rhizosphere microbes (Zobiole et al. 2011).

Control of *P. cinnamomi* by phosphite spray was no greater than the control seen in plants treated only with avocado orchard mulch. Metalaxyl had a greater detrimental effect on overall growth specially shoot growth of non-inoculated plants than phosphite. It has been observed that the ability of avocado trees to resist *P. cinnamomi* may increase with the age of an orchard (Develey□Riviere and Galiana 2007), and with the build-up of soil organic matter from leaf litter (Schadler et al. 2010). Leaf litter together with organic matter in particular well decomposed chick manure raised microbial activity and soil pH which in return significantly reduced the *P. cinnamomi* survival (Aryantha et al. 2000; Konam and Guest 2002). As soil microflora were not studied, it is possible that the proprietary microbial soil additives increased soil microflora populations, but they had no effect on *P. cinnamomi* beyond those provided by avocado orchard mulch. The avocado orchard mulch may either contain beneficial microbial consortia, and/or possess properties that enhance their growth.

## Conclusions

Although the experiment attempted to simulate the conditions of an orchard, it is difficult to exactly replicate orchard conditions in a container trial in glasshouse, and to be sure that the soil additives will have similar effects in the field. Temperature regimes, water retention, aeration, and surface evaporation are factors that might have different effects on abiotic conditions and thus microbiota in pots and in the field (Poorter et al. 2016). In particular, the microbes included in the probiotics may not multiply and persist as well under container conditions as in the field.

Control methods that reduce the use of chemicals or avoid the build-up of resistance to phosphite or fungicides are highly desirable for commercial horticulture. In the current study, the application of a number of soil additives to containerised avocado plants under glasshouse conditions indicated that avocado orchard mulch was beneficial in reducing Phytophthora root damage. Further control may be possible through the application of a silicon based mineral mulch. Probiotic conditioners did not improve disease control. The silicon content of the mineral mulch may enhance the plants defence mechanisms against the pathogen. Future experiments will investigate the effect of these treatments on the soil microbiome. It was also shown that application of glyphosate for weed control damages avocado roots. If chemical control of *Phytophthora* is necessary, phosphite is better if applied at recommended rates (Gilardi et al. 2020) than metalaxyl as it has less detrimental effects on overall plant growth. Phytophthora resistant isolates developed faster in metalaxyl than in phosphite treated plants, as the former chemical has a targeted mode of action (Browne and Viveros 2005), whilst phosphite does not (McDonald et al. 2001;

Browne and Viveros 2005). Under glasshouse conditions, none of the soil additives improved growth of potted avocado plants, or were more effective against *P. cinnamomi* than treatments simulating current silvicultural methods (application of chicken manure, wood mulch and retention of leaf litter) used by avocado growers in their orchards. However, it is important to conduct follow up trials by using these products in the field to confirm the findings of this study.

## Acknowledgement

This research was supported by Murdoch University Scholarship and HIA (Horticulture Innovation Australia) project AV10067. We thank Russel Delroy from Delroy farms for providing mulch from his orchard, Kay Howard for help with the preparation of the manuscript, Ziad Mekkawi, Mohemmad Bhaidhani, Ian McKernan and Bill Dunstan for assistance in the glasshouse, and Rajah Belhaj for lab work, Chris shaw and Peter Thomson for providing guidance in statistical analysis. We also thank Geoleaks Solutions, Mineral mulch and Eco Growth for providing commercial soil additives.

